# Massively calcified endosomal death (MCED) of endothelial cells

**DOI:** 10.1101/007112

**Authors:** L. Weisenthal

## Abstract

We have discovered a novel and specific mechanism of endothelial cell death. We refer to this novel death mechanism as massively calcified endosomal death, or MCED. Exposure of endothelial cells to non-specific toxins or other physical stresses induces death by traditional apoptotic and non-apoptotic mechanisms, common to most different types of cells. In contrast, exposure of endothelial cells (but not other types of nucleated cells) to specific insults, such as oxidized pathogenic lipids (e.g. 7-ketocholesterol) or agents with known anti-angiogenic activity (e.g. bevacizumab, certain tyrosine kinase inhibitors, etc.) triggers cell death via a novel pathway, which involves the formation of massively calcified endosomes, which, in turn, escape from the dying endothelial cells as massively calcified exosomes. These endosomes/exosomes appear capable of provoking an inflammatory response, characterized by physical association of calcified microparticles with inflammatory cells (monocytes, lymphocytes, neutrophils) with resulting increased release of an inflammatory mediator (TNF) into the culture medium. Traditional media for the culture of endothelial cells are profoundly inhibitory to MCED, as are some mammalian sera and many human sera, explaining why MCED had not been previously discovered and reported. The present discovery of MCED was accidental, resulting from work with primary cultures of fresh human tumor cell clusters, which invariably contain microcapillary cells. Our culture media are optimized for the tumor cells and not for the endothelial cells and, thus, are permissive of MCED. I propose MCED as the central mechanism underlying both intimal calcification and vascular inflammation in atherosclerosis.

## Introduction

Two major hallmarks of atherosclerotic vascular disease (ASVD) are calcification and inflammation. According to a current review^1^ “the mechanisms of intimal calcification remain poorly understood in humans.” According to another recent review^2^ “it is still unclear precisely what induces the inflammatory response.”

## Methods and results

Figure 1 shows human umbilical vein endothelial cells (HUVEC) which were incubated for 24 hours with 7-ketocholesterol, in RPMI-1640 medium with 20% fetal calf serum (screened for the absence of MCED inhibitors^3^) in 96 well plates, then 3% Fast Green dye in 0.9% NaCl was added and cells were sedimented onto microscope slides with a Cytospin centrifuge. Living cells (with intact cell membranes) remain clear (unstained). Non-specifically dead cells stain pale green, from having taken up Fast Green dye through incompetent cell membranes. Massively calcified endosomes stain densely refractile green (if counterstained with dyes such as hematoxylin and especially alizarin, these endosomes stain as vividly refractile, confluent “lakes” (their appearance as spherical endosomes is lost, in the process of interaction between calcium and the calcium stain, see Figure 4). In figure 1, the process of calcified endosome formation is at an early stage and the endothelial cell has not yet lost membrane integrity to allow general entry of Fast Green. It should be noted that stripping VEGF from the culture medium with bevacizumab also produces massively calcified microparticles (endosomes and exosomes), as in the case of exposure of HUVEC to 7-ketocholesterol (see Figure 4). However, exposure of HUVEC to a wide array of non-specifically toxic drugs does not produce this calcification (data not shown; but shown in reference^4^).

**Figure 1.**
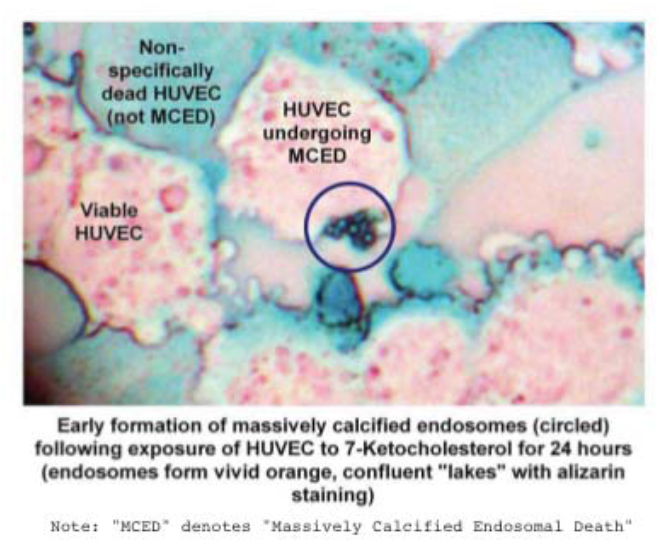

Figure 2 shows HUVEC which were incubated as described in Figure 1, but after 48 hours. In figure 2, the process of calcified endosome formation is at an advanced stage and the endothelial cell has lost membrane integrity to allow general entry of Fast Green.

**Figure 2.**
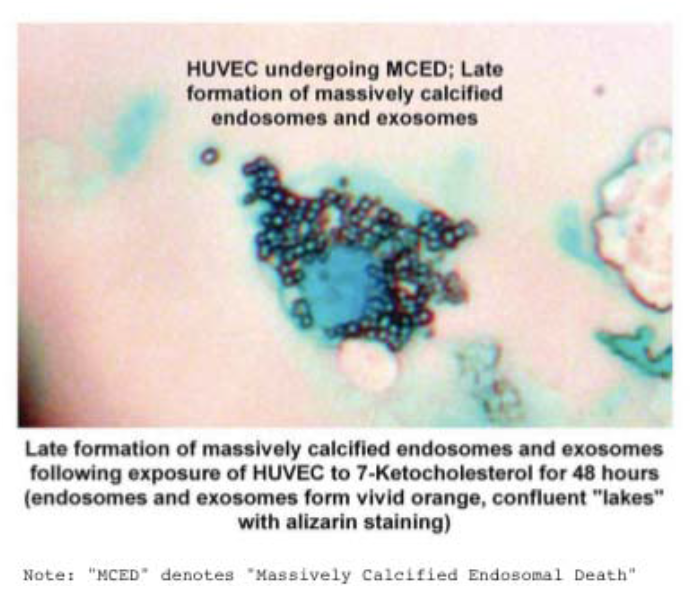

Figure 3 shows HUVEC which were incubated as described in Figure 2. In Figure 3, the process of calcified endosome formation is at a very advanced stage.

**Figure 3.**
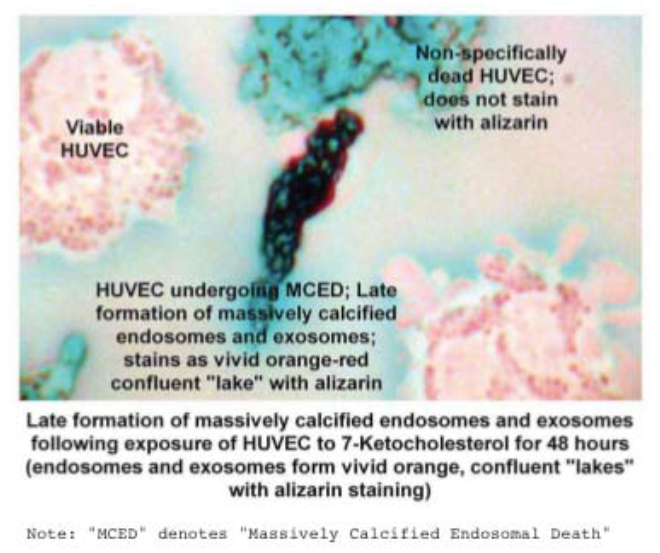

Figure 4 shows HUVEC which have been cultured for 96 hours in either (1) physiological saline vehicle, (2) doxorubicin 12 μg/ml, or (3) bevacizumab 2.5 mg/ml, a concentration which strips VEGF from the culture medium to undetectable levels^5^. It can be seen that exposure to doxorubicin produces only non-specific cell death, which does not involve the generation of massively calcified microparticles, while exposure to bevacizumab (or 7-ketocholesterol, Figures 1 - 3) produces dead cells permeated with massively calcified endosomes.

**Figure 4.**
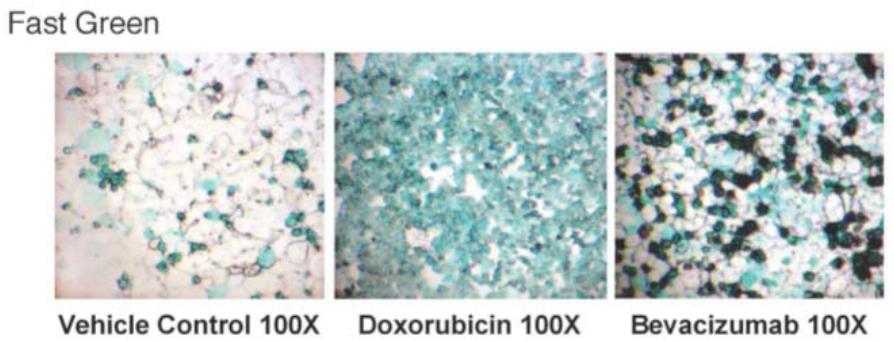
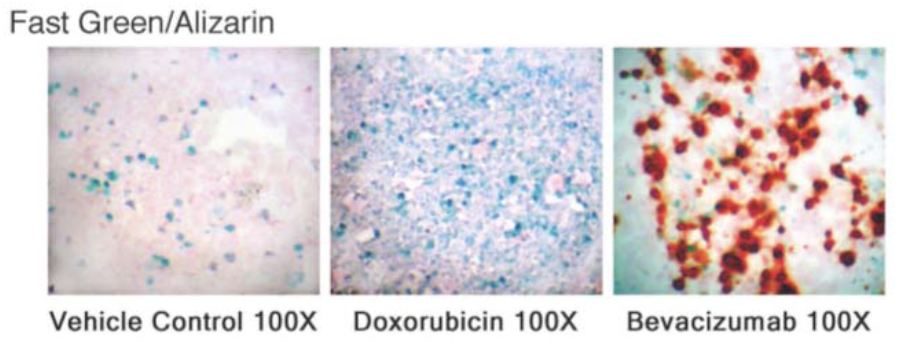
Living endothelial cells are clear (unstained); *non-specifically dead* endothelial cells — e.g. with cell death produced by exposure to the non-specific cytotoxic drug doxorubicin — stain pale green; *specifically dead* endothelial cells — e.g. produced by exposure to oxidized lipids or to specific anti-angiogenic drugs, such as bevacizumab in the present example — stain a highly refractile deep green. Upon counterstaining with Alizarin Red S — a specific stain for calcium — only the “specifically” dead endothelial cells show the presence of massive calcium.

Figure 5 shows that autologous calcified microparticles are capable of provoking an inflammatory response from autologous leukocytes incubated in autologous serum. Peripheral blood cells were incubated with bevacizumab (2.5 mg/ml, a concentration which effectively strips VEGF from the culture medium5) for 96 hours, as previously described5 to generate calcified microparticles. These calcified microparticles were then purified via centrifugation through Ficoll-diatrizoate, and then these calcified microparticles were incubated with freshly isolated autologous peripheral blood leukocytes. Cell culture supernatants were removed at 1 hour, 24 hours, 48 hours, and 72 hours and assayed for tumor necrosis factor (TNF) concentration via commercial ELISA, with paired comparisons between vehicle control-incubated and microparticle-incubated inflammatory cells. The accelerated release of TNF by peripheral blood cells in the presence of the calcified microparticles suggests that the calcified microparticles are capable of producing an inflammatory response.

**Figure 5.**
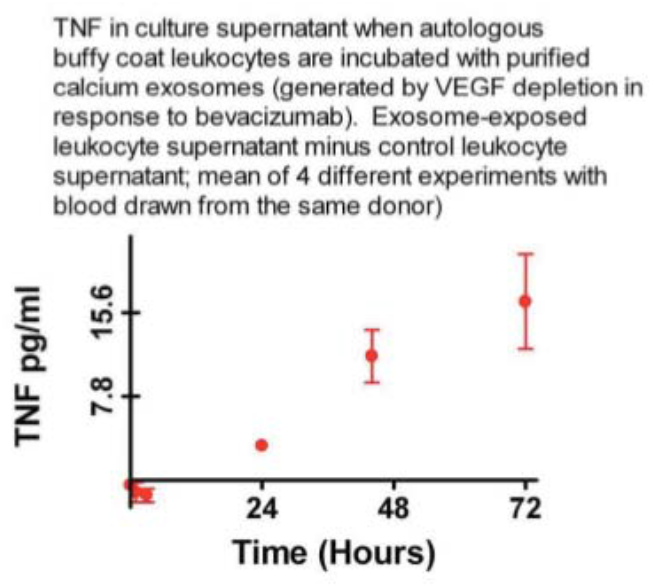

Figure 6 provides further evidence that autologous calcified microparticles are capable of provoking an inflammatory response from autologous leukocytes incubated in autologous serum. Peripheral blood cells were incubated with bevacizumab, in the same experiments as described in the legend for figure 4 to generate calcified microparticles. These calcified microparticles were then purified via centrifugation through Ficoll-diatrizoate, and then these calcified microparticles were incubated with freshly isolated autologous peripheral blood leukocytes. After 24 hours, these cells were stained with Fast Green, sedimented onto Cytospin slides, and counterstained with Wright-Giemsa. Under these conditions, the calcified microparticles stain a densely refractile blue-black, as previously described^5^. These experiments further suggest that autologous calcified microparticles are capable of provoking an inflammatory response from autologous leukocytes, as evidenced by physical interactions of inflammatory cells (lymphocytes, neutrophils, monocytes) with the calcified microparticles.

**Figure 6.**
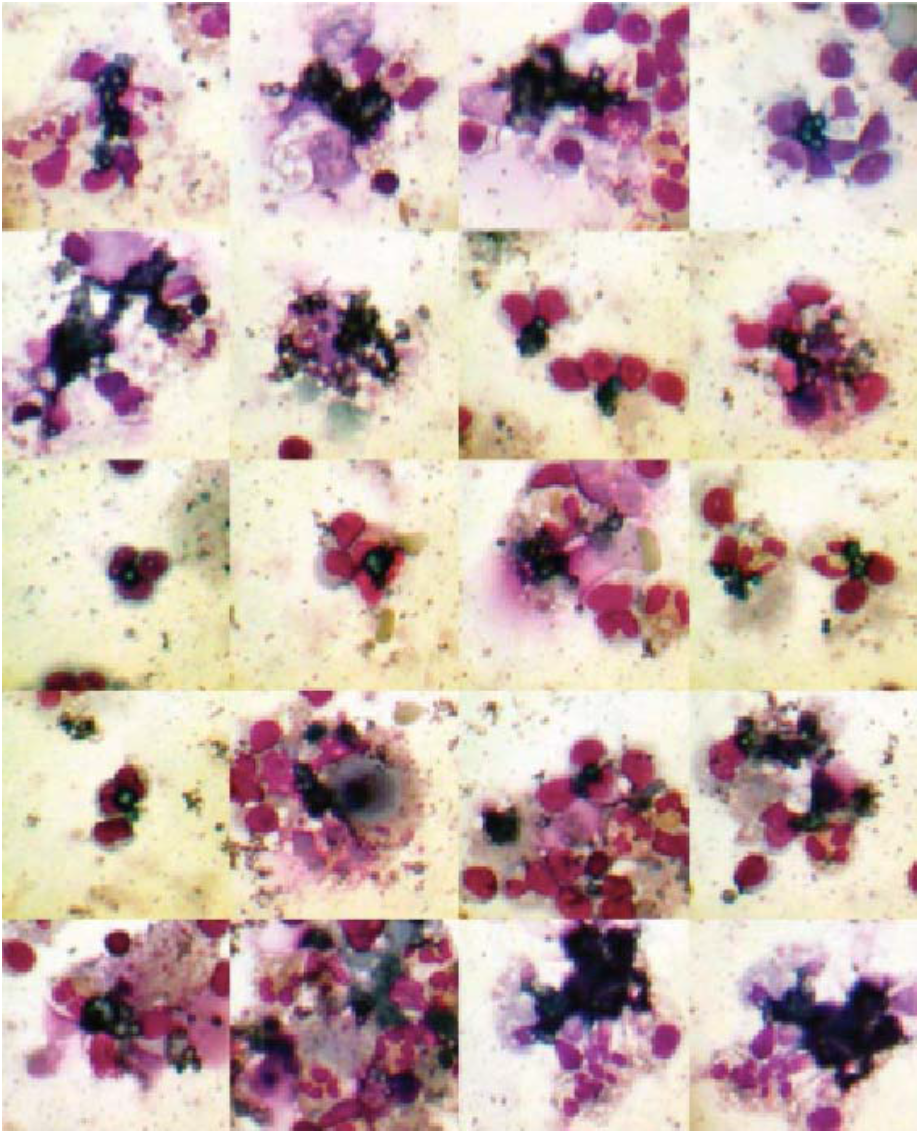

## Discussion

The present works shows that exposure of HUVEC to 7-ketocholesterol or to a concentration of bevacizumab which effectively strips VEGF from the culture medium generates massively calcified endosomes, which are released from the disintegrating endothelial cells as massively calcified exosomes. The present work also provides preliminary evidence that massively calcified microparticles are capable of generating an inflammatory response, as evidenced by physical association of autologous inflammatory cells (monocytes, neutrophils, and lymphocytes) with the calcified microparticles and accelerated of TNF into the culture medium when the microparticles are incubated with autologous peripheral blood leukocytes in the presence of autologous serum.

I propose massively calcified endosomal death (MCED) of endothelial cells as the underlying mechanism of both intimal calcification and inflammation in atherosclerotic vascular disease. I previously reported^3^ that a culture medium optimized for maintenance of endothelial cells completely blocks massive calcification of endothelial cells. I believe that this is the central reason why MCED was never previously discovered/described. The present discovery of MCED was entirely serendipitous, as this was discovered in experiments with primary cultures of three dimensional tumor microclusters, in which media optimized for endothelial cells was not utilized^5^.

